# Complex population dynamics in a spatial microbial ecosystem with *Physarum polycephalum*

**DOI:** 10.1101/2021.03.14.435300

**Authors:** Leo Epstein, Zeth Dubois, Jessica Smith, Yunha Lee, Kyle Harrington

## Abstract

This research addresses the interactions between the unicellular slime mold *Physarum polycephalum* and a red yeast in a spatial ecosystem over week-long imaging experiments. An inverse relationship between the growth rates of both species is shown, where *P. polycephalum* has positive growth when the red yeast has a negative growth rate and vice versa. The data also captures successional/oscillatory dynamics between both species. An advanced image analysis methodology for semantic segmentation is used to quantify population density over time, for all components of the ecosystem. We suggest that *P. polycephalum* is capable of exhibiting a sustainable feeding strategy by depositing a nutritive slime trail, allowing yeast to serve as a periodic food source. This opens a new direction of *P. polycephalum* research, where the population dynamics of spatial ecosystems can be readily quantified and complex ecological dynamics can be studied.

## Introduction

Experimental studies of microbial ecosystems have often yield insights into the relationships between the population dynamics of species. Futhermore, successional dynamics between species are often studied to understand the stability and sustainability of ecosystems, such as cyclic dynamics of ecosystems with rapidly evolving species (Yoshida et al., 2003), and stability of cooperativity in eco-evolutionary systems (Sanchez and Gore, 2013). This has led to suggestions of ecological mechanisms based on phenotypic dynamics (Harrington and Sanchez, 2014). However, experimental studies of microbial ecosystems generally focus on well-mixed systems, where quantitative measurements of population density can be directly acquired with spectrophotometery or flow cytometry. These types of measurements do not work for spatial ecosystems where spatial heterogeneity significantly influence measurements. In this work, we leverage experimental methods inspired by the Lifespan Machine (Stroustrup et al., 2013) to study a microbial ecosystem composed of *Physarum polycephalum* and a red yeast for week-long experiments. Our measurements capture the complete ecosystem with its original spatial structure using an advanced image analysis pipeline to quantify population dynamics. Through this approach we find an inverse growth relationship between *P. polycephalum* and a red yeast, suggesting that *P. polycephalum* acts as a predator and a red yeast serves as a prey. Additionally, we show multiple examples of successional population dynamics between the two species. This opens new directions for the study of *P. polycephalum* in the context of ecological dynamics.

### Background on *Physarum poylcephalum*

We outline some relevant background on *Physarum polycephalum* by discussing key mechanisms and behaviors of this unique model organism. We begin by describing morphological features of the organism that underlie its capabilities for acquiring and metabolizing food, exploring its environment, and adapting its morphology. We then discuss previous work on the interaction of *P. polycephalum* with microorganisms. Finally, we highlight recent work on external/stigmeric memory in *P. polycephalum*.

### Morphology

The peristaltic contractions of *P. polycephalum*’s vasculature has been measured and used to approximate information flow in the cytosol (Ray et al., 2019). This work suggests that some signal in the cytoplasm elicits a morphogenic response or performs a computation. By measuring peristaltic tubule contractions, they identify how *P. polycephalum* organizes fluid flow such that it can make a binary nutritional choice. In these experiments, a plasmodial tubule is grown between two nutritionally distinct food sources. The authors hypothesize that a solute contained within *P. polycephalum* cytoplasm contains some sort of signal that *P. polycephalum* uses to coordinate contraction across its vasculature. The dynamics of information transfer across the tubule are measured by calculating the rates of contraction across the length of the tube over time. Rates of contraction across the tubule are lower near to the higher quality food source and higher near to the lower quality food source. Ray et al. (2019) conclude that a signal is propagated from the end of the tube with a low-quality food source to the end of the tube with a high-quality food source. The fluid flow carries a signal that over some time horizon directs *P. polycephalum* movement toward the food source. Just as *P. polycephalum* distributes biological signals within its body to direct fluid flow, *P. polycephalum* may also leave behind signals in the environment.

There has been extensive work which has shown that *P. polycephalum* can perform embodied computation (Adamatzky et al., 2008). *P. polycephalum* can change its environment to more efficiently avoid negative stimuli and forage for food (Boussard et al., 2019). *P. polycephalum* deposits a slime sheath while migrating (Reid et al., 2012). This slime sheath, like many other cues in the environment (incline, light, heat, nutrition source, microorganisms, etc.) are detected by *P. polycephalum* and influence its behavior. These stimuli direct *P. polycephalum* migration and more generally morphogenesis (Vogel et al., 2016). In (Reid et al., 2012), *P. polycephalum* were placed on an agar plate in which a food source was blocked by an obstacle. In one condition the agar was coated with slime-sheath, in the other condition the agar was untreated. Across these two conditions, the time taken to reach the food source was measured. The *P. polycephalum* placed on a plate coated in slime-sheath took longer to navigate to the food than its counterpart in the untreated environment; in a coated environment the slime-sheath was a useless signal. In the untreated environment, by avoiding the slime-sheath *P. polycephalum* was able to navigate to food more efficiently. By modifying its environment *P. polycephalum* can better undertake complex tasks such as foraging. This observation clearly shows that biochemical signals in the environment interact with fluid flow to shape *P. polycephalum* morphogenesis.

### Interaction with microorganisms

With regards to the behavioral ecology of *P. polycephalum*, we test hypotheses about the ways in which *P. polycephalum* may interact with other microorganisms. We hypothesize that a red yeast eats the *P. polycephalum* slime sheath. This type of public good dynamic is common in Nature. Bacteria (in our case a protist *P. polycephalum*) produce metabolites that invite other microorganisms to grow near them (Nadell et al., 2016, 2008). *P. polycephalum* have been documented living in and under rotting logs; rich ecosystems; hotbeds of public good production and consumption. Interrogating how *P. polycephalum* interacts with a red yeast may help us contextualize many observations made about (fungi)taxis (movement towards fungal food sources) behavior in the lab. Working with image data we may characterize the spatiotemporal dynamics of an interspecies interaction. In our study we borrow heavily from the lexicon and methodology of ecology. Our experimental system is fertile for ecologists who would like to understand patchy unmixed interactions more generally. In many cases the dynamics of studied ecological systems work themselves out over times scales on the order of years, or occur on the spatial scale of biofilms. *P. polycephalum* and yeast provide a laboratory ecology that has dynamics that work themselves out over the course of approximately one week and can be observed without a microscope. It is also the case that *P. polycephalum* and the order *Mxyomycetes* play an important role in the ecosystems that they inhabit (as microvores) yet observations about their behavioral ecology are few and far between (Ing, 1994). Even in the existing *P. polycephalum* fungivory literature, the spatiotemporal dynamics of *P. polycephalum* – microorganism interactions have not been addressed in great detail.

Fungivorous and bacterivorous behavior has been observed across the class Mycetozoa (including *P. polycephalum*) (Chapman and Coote, 1983; Cohen, 1939). Both studies observe that *P. polycephalum* and other acellular slime molds such as *Badmania* grow much more vigorously when they are placed in an environment with yeast and other microorganisms. Others have observed that plasmodia growing in a two-member culture survive longer than they otherwise would and demonstrate a deepened behavioral repertoire (more dynamic growth) (Cohen, 1939; Chapman and Coote, 1983). Cohen (1939) hypothesizes that *P. polycephalum* grown in a two-member culture consumes the yeast while the yeast breaks down the complex carbohydrates locked up in the oats. In the context of our experiments, *P. polycephalum* may either utilize a nutrient-poor ring of oats, or invest energy in foraging for yeast. While *P. polycephalum* certainly consumes a red yeast, the time and tempo of this behaviour has not been characterized. *P. polycephalum* and yeast could have a predator-prey relationship, or the deposited slime sheath could introduce a successional dynamic into this bipartite ecological assemblage. A characterization of the interactions between *P. polycephalum* and red yeast may contribute to the sparse, but interesting, literature on *P. polycephalum’s* behavioral ecology.

In addition to predator-prey interactions with microorganisms *P. polycephalum* has been shown to enter into symbiotic relationships with other protists. Lazo (1961) has shown that, across many different *Myxomycete* (the class of organisms in which *P. polycephalum* belongs) – green algae species pairs, a symbiosis can arise. This work quantified symbiosis by measuring the color of the *Myxomycete* across time. As algae grows in the endoplasm the *Myxomycete* turns a shade of green (Lazo, 1961). Gastrich and Anderson (2002) observed that after several days albino *P. polycephalum* plasmodia would take on a green color. Gastrich and Anderson (2002) have shown that growing an albino variant of *P. polycephalum* in association with the photosynthetic protist, *Chlorella pyrenoidosa* increases the longevity of *P. polycephalum* from 5 to 10 days to up to a month. They assess symbiosis by measuring the green coloring of the albino *P. polycephalum*. They find it takes approximately one week for the plasmodia to turn green. Gastrich and Anderson (2002) use a TEM to perform ultra-structure analysis. They find the *Chlorella* within membrane compartments of *P. polycephalum*. The authors hypothesize that *Chlorella* can provide *P. polycephalum* with nutrients it produced through photosynthesis. *Chlorella’s* green pigment may protect the albino slime mold from harmful illumination. *P. polycephalum*, as well as other Mycetozoa, may form intimate beneficial relationships with many different types of microorganisms. To the best of our knowledge, others have not attempted to quantify the spatiotemporal dynamics of these sustained and unique types of interactions.

### External memory

Kataoka and Nakamori (2020) investigates the slime sheaths that many Myxomycete species deposit as they migrate. They demonstrate that different species of beetles in the genus Collembola may consume plasmodia, or the deposited slime sheath, exclusively. The deposited slime acts as externalized memory for P. polycephalum as well as a source of nutrients for other organisms (Reid et al., 2013; Kataoka and Nakamori, 2020). The authors note that, in the case of the plasmodia consuming insect, the slime mold is more likely to fragment - a single plasmodia splits up into smaller fragments. By observing the behavior of P. polycephalum, as well as other Myxomycetes, the authors of these studies make interesting and novel observations about Myxomycetes morphogenesis and the behavioral ecology of P. polycephalum. In our work, we observe a red yeast consuming a slime sheath. The slime sheath allows P. polycephalum to navigate through its environment, interact with other conspecific plasmodia, and serve as a food source. Importantly the slime sheath provides a means for P. polycephalum to establish an ecological relationship with other organisms.

Recently papers have demonstrated that P. polycephalum may modify its environment so that it may better avoid obstacles and find regions that contain food. Reid et al. (2013) has shown that P. polycephalum can navigate environmental obstacles such as barriers around food, by depositing a slime sheath in areas that they visit. When P. polycephalum travels across its environment it leaves behind portions of its actin skeleton and other unidentified components. Reid et al. (2013) demonstrate that P. polycephalum tends to avoid areas where they have deposited slime sheathing. By avoiding previously explored areas plasmodia are more efficiently able to explore their environment and navigate around obstacles through a process of elimination (unproductive paths are entombed by P. polycephalum’s slime sheath). Briard et al. (2020) discusses the topic of external memory and its relationship to slime mold’s deposition of its extracellular slime sheath. The observations in (Briard et al., 2020) offer new insights into the role of slime sheaths. In this work, the authors demonstrate that not all slime sheaths serve as an avoidant signal to other conspecific/clonal plasmodia. The signal of the slime sheath is highly correlated to the stress state of the plasmodia. Healthy plasmodia leave trails that are attractive to other conspecifics. Stressed plasmodia leave repellant slime sheaths. The authors show that P. polycephalum, when given a choice between regions explored by stressed plasmodia, regions explored by healthy plasmodia, and unexplored regions, will most likely go to the region with slime sheath deposited by the healthy plasmodia.

Alim (2018) suggests that physical forces may feedback into biochemical reactions to trigger complex spatiotemporal dynamics in the morphogenetic processes. It is possible that a similar feedback loop could be created when P. polycephalum consumes the red yeast. Given that (Briard et al., 2020) shows that a well-nourished P. polycephalum deposits an attractive slime sheath, as the plasmodia migrate in the environment and consume yeast, it might tend to leave more attractive slime sheath nearby sites of yeast predation. This site of slime sheath deposition could be consumed by the red yeast and serve as a future attractive stimulus. If not all slime sheath is consumed by the red yeast, the sheath would attract the P. polycephalum to the yeast once again. It is tempting to hypothesize that this stigmergic slime sheath could serve as a substrate for sustained successional ecological interaction. As times goes on in the experiment and as more of the dish is coated with the slime sheath patterns of plasmodial migration and taxis may become increasingly correlated with the spatial arrangement of the slime sheath and canalized in a self reinforcing loop.

### Experimental pipeline

An experimental pipeline was designed to capture the morphogenesis of *P. polycephalum* and its interactions with a red yeast over the course of a week. Each experiment contained the same components and was arranged in the same way. On a non-nutrient agar plate 12 oats were arranged on the perimeter of the plate like numbers on the face of a clock with *P. polycephalum* in the center. The oat at the 6 o’clock position was innoculated with a red yeast. These oats were used as food source to prolong our observations of *P. polycephalum* and the red yeast. While some might say that oats interfere with our attempts to quantify *P. polycephalum* yeast interactions, papers have shown that even in their presence interactions between *P. polycephalum* and microorganisms occur and do so at a great rate (Cohen, 1939). Oats are inert and were consistently placed across all experiments, this consistency makes it unlikely that they confounded our observations of *P. polycephalum* fungitaxis. An imaging platform was created to acquire a timeseries of images of the ecosystem, where the imaging platform was designed to be low cost and easily extensible.

*P. polycephalum* was grown in 100mm polystyrene petri dishes on non nutrient agar atop a flatbed scanner (Canon Lide200). On top of the scanner a wooden template is placed with six spaces for polystyrene dishes. The open source SANE (Scanner Access Now Easy) tool was used for command line control of the scanners. A bash script using the CRON daemon was written to schedule scans at regular intervals (every 5 minutes). This script was extended to control a blue light, turning it on and off at controlled intervals. Instead of a regular petri dish lid, dishes are fitted with a blacked out funnel, its sloping walls prevent condensation from forming. The black finish prevents outside light from reaching the *P. polycephalum*. A fitting on top of the black funnel enabled mounting of RGB LEDs for controllable exposure of blue light. Another template was placed above the cones so they would not fall over or shift over the course of the experiment. CRON jobs, light management, and file management were run using a Raspberry Pi. BASH scripts, run on the Pi, coordinate apparatus control. Finally an additional cardboard enclosure is placed over the entire device to ensure further stability as well as protection from external light sources.

In our study, we observe *P. polycephalum* morphogenesis across the scale of days. The images we acquire to analyze *P. polycephalum* morphogenesis are taken using a flatbed scanner. A subset of images were segmented by hand using the tool Ilastik (Berg et al., 2019). These hand segmented images were combined to create a curated ground truth training set for a U-Net model (Ronneberger et al., 2015a). Included in the training corpus are plates with a large quantity of healthy plasmodia, plates with a large quantity of dead plasmodia, plates with a large quantity of yeast, and plates in which these three classes interface. Plates close together in time are also included. These close-in-time plate pairs were included to ensure greater frame-to-frame class consistency. The trained U-Net model performs well, but is not perfect. Measuring and understanding the dynamics of robustness in neural networks is a very active area of research that requires some mechanistic knowledge of the neural network learning process, which the field does not have. No optimal U-Net training regime is known to exist.

With our experimental pipeline we attempt to create a dataset that can be used to characterize *P. polycephalum* and red yeast interactions. We hypothesize that they interact in a way that can be captured across the time series of images that we produce. A predator-prey interaction is feasible between these two organisms, as *P. polycephalum* consumes the red yeast, red yeast consumes the slime sheath that *P. polycephalum* leaves behind, and finally the *P. polycephalum* migrates around the dish, leaving in its wake the slime sheath which can be consumed by the red yeast. We also expect that it may be difficult to consistently observe these interactions because spatial ecosystems are known to both be sensitive to initial configurations and have chaotic dynamics (Wakano et al., 2009). In the context of our study, *P. polycephalum* will not always be able to find yeast to eat, and yeast will not be able to grow on inert slime sheath.

### Quantifying Population Dynamics

When interactions occur and we quantify the population dynamics. The key challenge of analyzing our experiments is the image segmentation step. The experimental observations consist of 2,000 images for each timeseries, where each image has been acquired at a 5-minute interval. We developed an advanced image analysis procedure to assess the area covered by each species, leading to quantified population dynamics such as those shown in Figure 3. Code for the experiments is made freely available^1^.

### Semi-automated segmentation

Ilastik pixel segmentation was used to create and curate interactively labeled ground truth masks. Ilastik is an interactive random forest classifier used for image analysis (Berg et al., 2019). The workflow for Ilastik pixel segmentation is straightforward and efficient. Portions of an image are hand segmented by the user. These segmentation annotations are then used by the Ilastik classifier to train a random forest classifier that operates on a collection of filtered images to segment the remaining unclassified regions of the image. The user may add or remove annotations from the image used to further refine the Ilastik classifier. Iterative annotation is carried out until a satisfactory segmentation is produced by Ilastik. A segmentation is considered sufficient when a large portion of the protoplasm rich plasmodia are accurately segmented and its vascular morphology is well preserved. Holes in the plasmodial vasculature are well defined and blebbing across looks accurate. To create the annotations required to train the Ilastik classifier, portions of our images were labelled as one of the following classes: protoplasm rich *P. polycephalum*, protoplasm poor *P. polycephalum*, red yeast, oats, and background. In Figure 2, examples of these classes and their annotations are shown. The Ilastik feature filters that were used as input to the segmentation were color intensity with a sigma of 0.3, edge with a sigma of 0.7, and texture with a sigma of 0.7. Sigma refers to the width of the Gaussian kernel used to smooth the image before application of the feature filters. The three sigmas chosen for each filter were the smallest available. With smaller sigmas it is easier to produce a semantic segmentation that faithfully captures the delicate structure of the *P. polycephalum* vasculature and more faithfully captures finer detail in the semantic segmentation.

The segmentation annotations used to train the Ilastik classifier were constructed by finding circular regions (with a radius of 31 to 61 pixels) of the image that contained as many of the aforementioned classes, as possible and fully annotating them. By fully annotating these circles we can define the boundaries between each of the classes which may appear very similar. One such boundary would be between the red yeast and depleted protoplasm. Both of these classes have the same translucent white color, differing only in shape and texture. This approach to annotation advances our stated goal of by capturing the nuances of the *P. polycephalum* vasculature, accurately capturing the shape of objects from each class, the shape of the interface between classes, and the texture of the different features. Ilastik provides a straight-forward and natural user interface to carry out interactive segmentation with a powerful random forest backend.

### Deep learning-based segmentation

While the Ilastik segmentation is quite robust across an image, there are several weaknesses inherent in the method that make it difficult to use as the only segmentation tool in our study. For example, when more annotations are added to the classifier the run-time can increase from minutes to tens of minutes, even during the annotation process. Training a robust model is inhibited by memory and run-time constraints. We also found that the accuracy of Ilastik segmentations drops when used on different experiments and scanners. This change in performance may be associated with artifacts that are unique to the experiment: plasmodial thickness, vascular architecture, as well as the spatial organization of *P. polycephalum* and yeast interaction. To overcome these issues we interactively train several Ilastik classifiers on different experiments, and then collate their outputs into a training corpus for a neural network with a U-Net architecture (Ronneberger et al., 2015b). The U-Net architecture has become very popular in the field of biological image analysis recently due to good performance across many different semantic segmentation tasks. By taking segmentations from many different Ilastik classifiers we hope to create a more robust image segmentation method. Simply put, we are able to quickly train several Ilastik classifiers to accurately segment a single image. These images can have widely differing abundances of semantic segmentation labels as well as different brightnesses, luminosities and unique scanner artifacts. With a U-Net we unify these many random forests-produced results into a single robust deep learning-based model. Similar approaches for bootstrapping segmentation training data from Ilastik to U-Net have been shown to be successful (Pape et al., 2021).

### Data sorting

With these pixel-wise semantic segmentations we set out to measure biological processes associated with *P. polycephalum* morphogenesis. Each of the six plates across each scanner run were separated into a segmented time series with each plate annotated with the previously described semi-automated segmentation classes. To create a mask of the circular plate, we used our knowledge of the experimental set up. In each scanner image there is a light wooden frame which has holes for six polystyrene plates full of media. These polystyrene agar dishes are under a black cone that blocks *P. polycephalum* from outside illumination. These light-blocking cones create six black circles against a light wooden background. These six circles have a similar radius and color. By using a Hough circle detector with the radius, number and distance between the centers of these dark circles as hyper parameters we find the coordinates of each plate early on in the experiment and use these coordinates to split the scanned images into six images of each dish. The plates are stationary during the scanning process, allowing the coordinates from the Hough circle detector to be used for the entire time series. As each experiment has a slightly different placement of the wooden frame the Hough circle detector is run one time for each experimental run. A substantial amount of work was put into creating an imaging platform with which we could keep *P. polycephalum* healthy for an extended period of time, while also ensuring that the imaging quality would be sufficient to observe the ecosystem’s behavior.

### Detection interaction areas

We suppress frame-to-frame variability in segmentations by taking two adjacent frames and comparing them. Any pixel with the identity of *P. polycephalum* in both images is labeled *P. polycephalum* in the denoised new image. Pixel pairs with only a single *P. polycephalum* identity are labeled as background. This process results in fewer noisy pixels along class boundaries and fewer scanner artifacts, making it easier to identify short term trends. This filtering is carried across all time series. We are also interested in measuring morphogenesis over longer temporal horizons (more than a couple of frames). In this case other error suppression rules are used. Across fifty consecutive frames, pixel values are summed along the time axis and the median plasmodial age, variance, and mass are computed. Several frames at the beginning and the end of the fifty frame window are used to make binary masks based on presence or absence of pixels with the identity of protoplasm rich plasmodia. These masks are compared with a presence/absence mask of *P. polycephalum* across the rest of the temporal window. These comparisons give us some metric of *P. polycephalum* growth and retraction.

### Analysis and results

We present and discuss observations made over 5 sample ecosystems where we were able to capture the interaction between *P. polycephalum* and red yeast. The raw time series of population densities in interaction areas are shown in Figure 3 for both species. These samples have some common trends, where for the first part of the timeseries (approximately the first 750 timesteps) both species are in a growth phase and have minimal interactions, and at the end of the timeseries inverse growth rate relationships and oscilliatory dynamics can be observed. These samples are taken from 3 light-based environmental perturbations (no light: Figure 3a, constant light: Figure 3b, 50% randomized light exposure: Figures 3c, 3d, 3e) that were introduced to survey the robustness and potential controllability of the ecosystem. The challenge of consistently ensuring interactions between species across many experiments prevents us from making inferences about the impact of light exposure on ecological interactions. Therefore, we restrict our observations about the impact of light exposure on the ecological interactions to simply representing a source of environmental variation. We note that even in the presence of this source of environmental variation our observations about inverse growth rates and oscillatory dynamics is consistent.

### Population dynamics of *P. polycephalum* and a red yeast

The spatiotemporal dynamics of *P. polycephalum* and red yeast’s interactions are complex and difficult to capture. At the start of each trial *P. polycephalum* and the red yeast are placed in different areas of the dish. This means that there is usually significant time lag before the two organisms interact, if they interact. When *P. polycephalum* interacts with yeast, it consume it (see Figure 1). *P. polycephalum* may invest the energy it has harvested from yeast, as well as oats into many behaviors. The two behaviors which are most easily measured are growth and migration. Fitting these complex processes into a single time series is a challenging task. Migration and growth might obscure *P. polycephalum* and yeast interactions in a univariate time series of *P. polycephalum* mass. *P. polycephalum* and yeast interactions often take place in a very small portion of the dish. While we have gathered data of *P. polycephalum* sweeping over expansive lawns of a red yeast^2^ this is not the most common form of interaction between the two species. Red yeast takes a long time to grow and *P. polycephalum* motility is influenced by age and nutritional status. We found that a more robust way of measuring the interactions between *P. polycephalum* and yeast was to measure the regions in a dish where *P. polycephalum* and yeast both occur across the whole time series (see Figure 3). In doing so the observations are more likely to capture *P. polycephalum* and yeast interactions especially if there are sustained oscillations between the two species.

**Figure 1.**
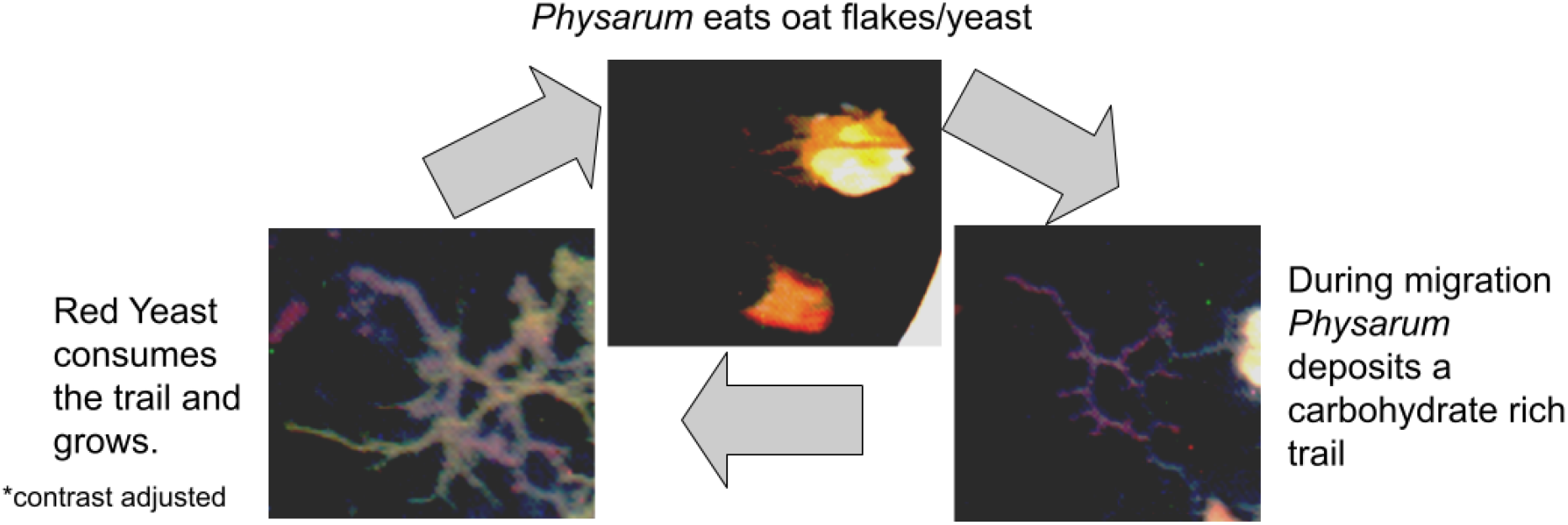
Diagram of the ecological dynamic in the experimental system.

**Figure 2.**
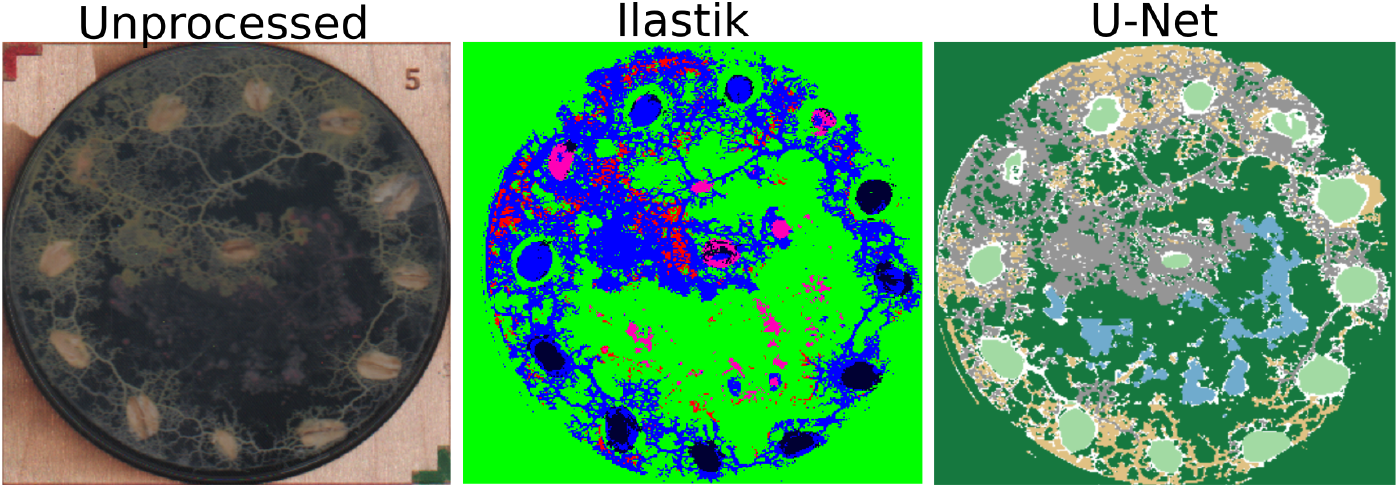
In the side by side comparison of the two segmentation strategies and the ground truth images we can clearly see that U-Net more faithfully captures the vascular architecture of *P. polycephalum*; protoplasm rich (labeled with blue and grey) and protoplasm poor (labeled with pink and gold), than its Ilastik counterpart. While neither segmentation algorithm can identify all of the yeast colonies (labeled in red and blue), U-Net captures many more than Ilastik.

**Figure 3.**
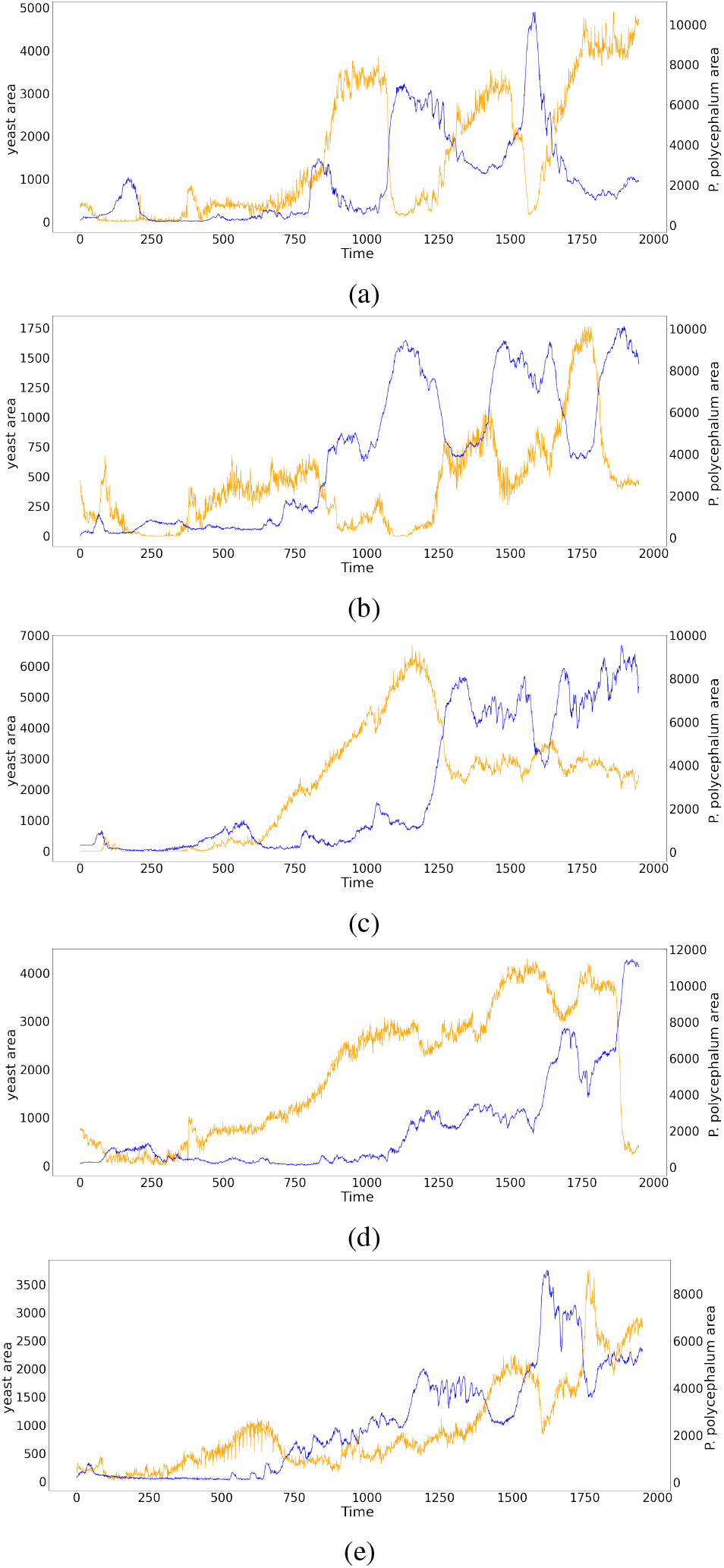
Timeseries of 5 ecosystems. Area (in pixels) of protoplasm rich *Physarum polycephalum* is shown in blue with values on the right-side axis. Area (in pixels) of red yeast is shown in yellow with values on the left-side axis. Three light conditions were used to survey the robustness and potential controllability of the ecosystem: no light exposure (3a), constant light exposure (3b), and randomized light (for 50% of the experiment; 3c-3e).

The growth rates of *P. polycephalum* and the red yeast have an inverse relationship. We show this through a phase plot that relates the change in normalized density of protoplasm rich *P. polycephalum* to the change in density of red yeast (see Figure 4). Densities are normalized to simplify comparison across samples. Here we show that there is a consistent correlation of inverse growth rates, where positive yeast growth rates correlate to negative growth rates of *P. polycephalum* and negative yeast growth rates correlate with positive growth rates of *P. polycephalum*. The *R*^2^ values support the trend, especially when considering the noisy aspects of the experiments, which include the stochasticity of the underlying biological processes, variation in images, and variation in segmentation outputs. This offers support for the hypothesis that *P. polycephalum* and red yeast have a predator-prey relationship.

**Figure 4.**
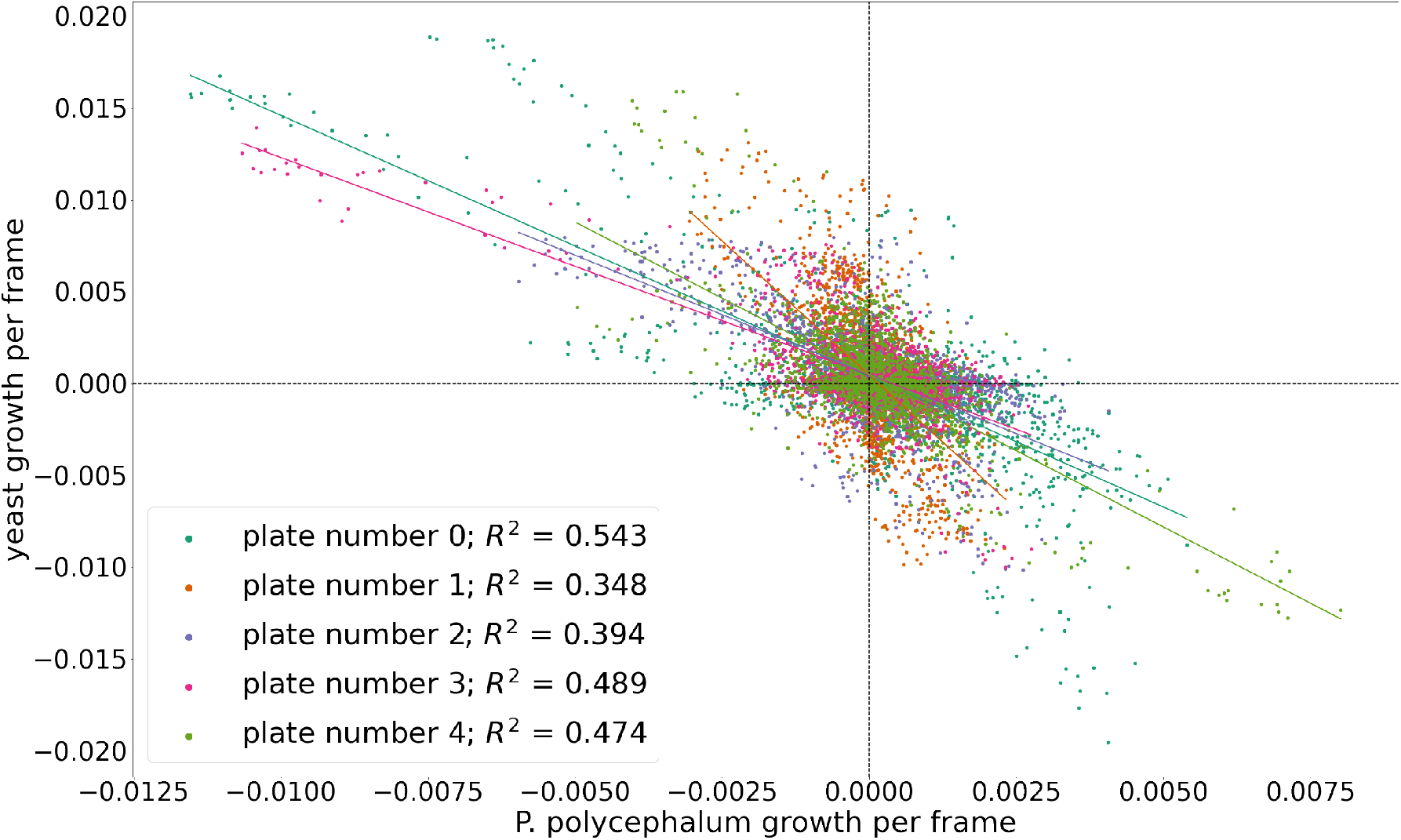
Phase plot of growth rates of *Physarum polycephalum* and red yeast at each timepoint for all timeseries presented.

If we now revisit the timeseries shown in Figure 3 we can address the oscillatory dynamics. These oscillations are most clearly shown in Figure 3a, where 3 cycles of the oscillation between *P. polycephalum* and yeast are shown. As opposed to idealized systems like the Lotka-Volterra mathematical model, our ecosystems are not capable of exhibiting steady state dynamics. The food sources that are available in the ecosystem will eventually be depleted. The spatial constraints of the ecosystem introduce local interactions that can exhibit chaotic dynamics which are even present in idealized mathematical models (Wakano et al., 2009). In Figure 3b, some oscillations are observed, but it is clear that the area covered by each species may operate on differing orders of magnitude. This is clear in the remaining samples shown in Figure 3. Nevertheless, we reiterate that there is a consistent correlation of inverse growth rates between the species, which is shown in Figure 4. The key to finding and quantifying this interaction in a way that is comparable across experiments was restricting the analysis to zones of interaction that leverages spatiotemporal information of both species.

## Conclusions

*P. polycephalum* has been shown to be fungivorous (Chapman and Coote, 1983) and others have engineered symbiotic relationships between *P. polycephalum* and other protists (Lazo, 1961; Gastrich and Anderson, 2002). In this work, we report an inverse growth relationship in an ecosystem composed of *P. polycephalum* and yeast, and present examples of oscillatory successional dynamics between the species. The question remains: what is the mechanism of this multispecies interaction? A recent paper by Kataoka and Nakamori (2020) mentions that a beetle may eat the slime sheath produced by the migrating *P. polycephalum*. The slime sheath is a source of nutrition for the beetle. The interactions between *P. polycephalum* and red yeast may be sustained by nutritive qualities of *P. polycephalum* slime sheath. As *P. polycephalum* leaves an area it deposits slime sheath. We have observed that the slime sheath is subsequently consumed by a growing colony of red yeast (see the supplemental video in footnote 2). This red yeast is then consumed by *P. polycephalum*. While additional experiments would need to be carried out to determine the exact nutritive properties of *P. polycephalum* slime sheath. The slime sheath provides a plausible explanation as to what drives this interaction. *P. polycephalum’s* slime sheath has also been shown to help clonal *P. polycephalum* plasmodia navigate their environment. Healthy plasmodia leave behind an attractive stimuli in their slime sheath and stressed Plasmodia leave behind a repellant stimuli in their slime sheath (Briard et al., 2020). We speculate that this slime sheath may have been consumed by microorganisms, just as it has been shown to be consumed by beetles. Our observation could add a new layer of complexity to the biology of *P. polycephalum* slime sheath. Following this observation we speculate that as *P. polycephalum* navigates its environment it occupies specific regions of a morphospace so that it may build a spatiotemporally optimal slime sheath to farm the most microorganisms. Significantly, we present quantitative observations, for the first time, of successional dynamics between *P. polycephalum* and another microorganism.

We were able to observe and report a new behavior in *P. polycephalum*. We were able to quantitatively measure population dynamics that suggest successional dynamics between *P. polycephalum* and red yeast. While *P. polycephalum* is often cited for its capacity to approximate shortest paths between different food sources, there are other facets of its behavior that warrant attention. Briard et al. (2020) shows that *P. polycephalum* slime sheath can provide an attractive or repulsive stimulus depending on the nutritive state of the depositing plasmodia. These can aid in navigation, helping *P. polycephalum* more robustly navigate its environment in search of food (Boussard et al., 2019). We suggest that this deposited slime sheath may also act as a food source for microorganisms. This would expand the importance of the slime sheath and extend our understanding of *P. polycephalum’s* role in ecological interactions. We have addressed key challenges in experimentally quantifying population dynamics in a spatial ecosystem composed on *P. polycephalum* and red yeast. As a whole, we have shown that there is an inverse relationship between the growth rates of *P. polycephalum* and a red yeast, and have demonstrated that this can lead to successional dynamics between the two species.

## Future Work

We suggest that extensions of this work should investigate ways to leverage the spatial configuration of the ecosystem to guide the ecology. The capacity to control the ecosystem with light was not clear. It may be particularly interesting to continue to investigate ways to maintain sustainable interactions between the species to observe population dynamics over longer time periods.

## Acknowledgements

Research reported in this publication was partially supported by the National Institute Of General Medical Sciences of the National Institutes of Health under Award Number P20GM104420. The content is solely the responsibility of the authors and does not necessarily represent the official views of the National Institutes of Health. This research was also supported by a Seed Grant from the University of Idaho Office of Research and Economic Development.

## Footnotes

1 Repository for the source code is here https://github.com/ComputationalAndPhysicalSystems/physarum_ecology

2 For an example of this compelling interaction, see the supplemental video available here https://youtu.be/IE7ZyzPW5Pc

